# Docking of peptides to GPCRs using a combination of CABS-dock with FlexPepDock refinement

**DOI:** 10.1101/2020.03.21.001396

**Authors:** Aleksandra E. Badaczewska-Dawid, Sebastian Kmiecik, Michał Koliński

**Affiliations:** Department of Chemistry, Iowa State University, 2415 Osborn Dr, Ames, IA 50011, USA; Faculty of Chemistry, Biological and Chemical Research Centre, University of Warsaw, 1 Pasteura St., 02-093 Warsaw, Poland; Bioinformatics Laboratory, Mossakowski Medical Research Centre, Polish Academy of Sciences, 5 Pawińskiego St., 02-106 Warsaw, Poland

**Keywords:** GPCR-peptide interaction, peptide docking, flexible docking, receptor-peptide complex, peptide drugs, model refinement

## Abstract

The structural description of peptide ligands bound to G protein-coupled receptors (GPCRs) is important for the discovery of new drugs and deeper understanding of the molecular mechanisms of life. Here we describe a three-stage protocol for the molecular docking of peptides to GPCRs using a set of different programs: (1) CABS-dock for docking fully flexible peptides; (2) PD2 method for the reconstruction of atomistic structures from C-alpha traces provided by CABS-dock and (3) Rosetta FlexPepDock for the refinement of protein-peptide complex structures and model scoring. We evaluated the proposed protocol on the set of 7 different GPCR-peptide complexes (including one containing a cyclic peptide) for which crystallographic structures are available. We show that CABS-dock produces high resolution models in the sets of top-scored models. These sets of models, after reconstruction to all-atom representation, can be further improved by Rosetta high-resolution refinement and/or minimization, leading in most of the cases to sub-Angstrom accuracy in terms of interface RMSD measure.

## Introduction

In the last two decades, peptides have gained a significant interest as therapeutic agents [1]. As demonstrated in many drug design studies, peptides can be useful as leading molecules and an alternative to small-molecule and biological therapeutics [1]. GPCRs are the largest and the most diverse family of membrane receptor proteins and are key drug targets [2]. It has been estimated that among the 826 human GPCRs, 118 can bind endogenous peptides and 30 are targeted by approved drug molecules [3]. Due to the difficulties in crystallization of GPCRs [4] only a few experimental structures of the peptide-bound receptor are now available [5]. In this context, there is growing demand for the development of efficient tools for the accurate computational prediction of GPCR-peptide complexes.

Due to the large interest in peptide therapeutics, many new protein-peptide docking techniques have been developed recently [6]. These may be divided into three categories: template-based docking, local docking and global docking tools [6]. Template-based docking uses structural data from analogous protein-peptide complexes. In the local docking the search for peptide-bound conformation is limited to the vicinity of the expected binding site. Finally the global docking methods perform search over the entire receptor surface.

Docking molecules to membrane proteins is a challenging task due to their large hydrophobic surface and the effect of the membrane environment that should be considered. There are few available methods that enable docking small molecules to GPCRs [7–10]. These may also be applied to short peptides, up to 5 residues [11]. Docking of larger peptides requires tools dedicated to flexible peptide docking, protein-protein docking, or modeling protocols tailored to particular GPCR-peptide systems. The examples include template-based modeling using Rosetta [12], manual docking guided by NMR data [13], CABS-dock docking followed by selection of plausible models [14], application of the hybrid molecular modeling protocol that integrates heterogeneous experimental data with force field-based calculations [15], application of ZDOCK and RDOCK tools for protein-protein docking [16] and GalaxyPepDock [17]. Nevertheless, the performance of neither of these methods was evaluated by prediction of a larger set of different GPCR-peptide complexes.

The GPCRs ligand binding site is located in the cavity formed by the bundle of seven alpha-helices. Larger ligands, like peptides, can also interact with extracellular receptor fragments (three extracellular loops and N-terminal domain) [18]. An effective procedure for GPCR-peptide docking should allow for full peptide flexibility, which can be crucial for the peptide deep penetration of the binding cavity and adaptation to the shape of the receptor extracellular surface. Moreover, all GPCRs show significant conformational flexibility and agonist binding is responsible for the stabilization of diverse receptor activation states [19]. Therefore, some receptor flexibility should be also accounted for during the docking procedure.

In this work, we present the modeling protocol dedicated for prediction of GPCR-peptide complexes that combines a coarse-grained CABS-dock global docking tool [20] with PD2 [21] reconstruction of the backbone and C-beta atoms positions and high-resolution Rosetta FlexPepDock refinement and scoring [22, 23] of protein-peptide complexes. During CABS-dock docking simulation the search for the receptor-peptide interaction interface is limited to the broad area of the extracellular parts of GPCRs. The proposed method was evaluated during the test prediction of 7 different GPCR-peptide complexes for which the crystal structures are available. The results of these docking experiments demonstrate that the proposed procedure enables highly accurate prediction of peptide binding poses.

## Materials and methods

This section starts from the description of the benchmark dataset used during evaluation of the proposed protocol. Then we describe the three consecutive stages of the modeling pipeline that are also illustrated in Figure 1.

**Figure 1.**
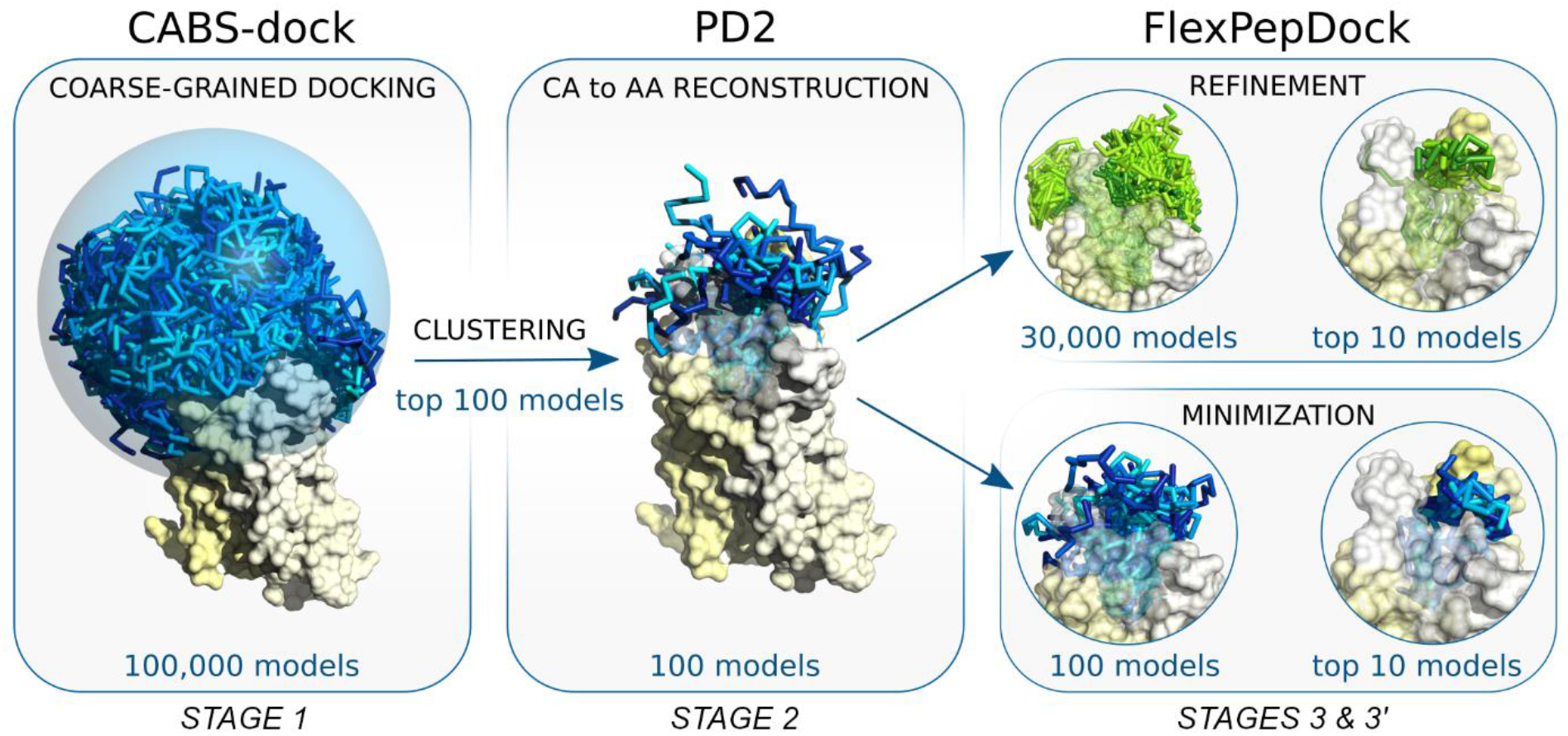
The pipeline used for docking peptides to GPCRs. The main stages of the docking procedure include: STAGE 1: Generation of a large number (100 K) of low energy receptor-peptide complex models resulting from 10 independent CABS-dock simulations; STAGE 2: Reconstruction to all-atom representation of 100 top selected models using PD2 [21]; STAGE 3: Model refinement and scoring using Rosetta FlexPepDock [22, 23].

### GPCR benchmark dataset

For preparation of the GPCR dataset used in this study, we examined all crystal structures of receptor-peptide and -peptidomimetic complexes form the largest Class A of GPCRs [24]. From the set of 9 identified crystal structures 2 were rejected due to the content of additional non-standard amino acid residues in the peptide ligand chains, which could not be properly parametrized according to the CABS-dock coarse-grained model. Structures of 7 different receptors were extracted from crystal structure files (PDB ID; reference; name; resolution): 3OE0 [25] - CXCR4 receptor of 2.9 Å; 4GRV [26] - NTS_1_ receptor of 2.8 Å; 5GLH [27] – ET_B_ receptor of 2.8 Å; 5XJM [28] - AT_2_ receptor of 3.2 Å; 6C1R [29] - C5a_1_ receptor of 2.2 Å; 6DDF [30] - μ-opioid receptor of 3.5 Å; 6OS9 [31] - NTSR_1_ receptor of 3.0 Å and used in the docking procedure. Peptide length varied from 5 to 21 amino acid residues. In the sequences of 4 docked peptides non-standard amino acid residues and/or D-amino acids were replaced by standard amino acid residues based on structure similarity criteria. This step was necessary because the CABS model (the CABS-dock simulation engine) supports only 20 standard L-amino acid residues. For sequences of all docked peptide ligands see Supplementary Table S1.

### STAGE 1: Molecular docking using CABS-dock

CABS-dock is a well-established method for flexible protein-peptide docking and its current status has been summarized in the recent review [32]. CABS-dock uses a very efficient simulation approach, the CABS coarse-grained protein model (its broad applications in protein modeling, protein structure prediction, simulation of protein flexibility and disordered states have been recently summarized in the reviews [33–35]. CABS-dock has been first introduced as a web server [36–38] and successfully applied to modeling large-scale conformation changes of protein receptor during peptide binding [39], protein-protein docking [38], peptide docking using sparse information on protein-peptide residue-residue contacts [40] and modeling the cleavage events occurring during proteolytic peptide degradation [41]. Recently, CABS-dock has been made available as a standalone application [20], which contains many features and extensions for advanced users. The repository of the CABS-dock package, which includes tutorials and description of commands, is available at https://bitbucket.org/lcbio/cabsdock/.

In this work, we used the CABS-dock standalone application [20] with the following input information (the structural data including peptide sequence information and restraint parameters used for CABS-dock input preparation for each of the 7 modeled systems are presented in Supplementary Table S1):

- amino acid sequences of docked peptides,
- coordinates of C-alpha atoms of selected receptor proteins (see section GPCR benchmark dataset),
- description of the peptide sampling space covering all GPCR fragments which may interact with bonded peptides (in the form of long-range distance restraints between receptor and peptide C-alpha atom pairs),
- information on internal disulfide bonds within peptide molecules (in the form of short-range distance restraints imposed on C-alpha atom pairs of particular cysteine residues),
- information on the cyclic conformation of one of peptide molecules (in the form of short-range distance restraint imposed on the C-alpha atom pair of selected residues).

The CABS-dock method has been designed for global docking simulations in which peptides are allowed to interact with the entire protein surface. In the case of GPCRs a large hydrophobic surface is responsible for positioning the protein in the membrane and this part of the protein does not interact with bound ligands. Therefore, in the docking simulations, we limited the peptide conformational sampling space to the broad neighborhood of the extracellular receptor domain including three extracellular loops and the receptor binding site located in the cavity formed by a seven-alpha-helical bundle (see Figure 1). Technically, the peptide sampling space was limited to a large sphere; with a radius of 25 Å or 30 Å, depending on the peptide size, using properly adjusted distance restraints (technical details describing the spherical docking space are provided in the Supplementary section ‘Technical details of the proposed workflow’). The restraints did not have any influence on peptide motion or the scoring within the sphere radius. This crude but simple and effective approach prevented peptides from binding to the surface of the receptor trans-membrane and intracellular domains, that is, the GPCR regions that do not interact with bound ligands. Additionally, as the input we used the information regarding disulfide bonds within peptide molecules in two modeled systems: PDB ID: 5GLH and 3OE0 (using relatively strong distance restraints between pairs of C-alpha atoms of appropriate cysteine residues). In the 6C1R system distance restrains were also imposed between C-alpha atoms of Ala2 and Arg6 of the docked peptide to preserve its cyclic conformation.

For each of 7 studied GPCR-peptide complexes, we performed 10 independent CABS-dock simulations. The CABS-dock commands and PDB structure files used for the modeling of all the receptor-peptide system are provided in Supplementary Table S1 and Table S2. The command line for running single docking simulation for the 6DDF system is presented below:

~~~
~/CABSdock -s *100* -M -C -S -v *4* -i *6DDF_struc.pdb:R* -p *6DDF_struc.pdb:D*
--reference-pdb *6DDF_struc.pdb:R:D* --sc-rest-add *147:R 1:PEP 25.0 5.0*
--sc-rest-add *147:R 2:PEP 25.0 5.0* --sc-rest-add *147:R 3:PEP 25.0 5.0*
--sc-rest-add *147:R 4:PEP 25.0 5.0* --sc-rest-add *147:R 5:PEP 25.0 5.0*
~~~

Each of 10 independent CABS-dock simulation runs generated 10 top-scored models (10 top-scored models were selected from the set of 1000 lowest energy models using the structural clustering approach). Next, we merged the sets of the top 10 models from 10 simulation runs into a cumulative set of top 100 models. This model set was further used as the input in STAGE 2 of the modeling protocol for the reconstruction of atomistic structures.

### STAGE 2: Structure reconstruction using PD2

CABS-dock standalone outputs provide models of protein-peptide complexes in C-alpha or all-atom representation. All-atom models can be generated thanks to CABS-dock integration with the Modeller-based reconstruction procedure (as described in the CABS-dock standalone documentation). In this pipeline, however, we recommend using the PD2 tool, since it has been proven to be slightly more accurate and more convenient in our internal tests than the Modeller-based procedure. The PD2 method enables fast and accurate reconstruction of the protein main chain and C-beta from C-alpha trace or partial backbone structures [21]. What’s important, PD2 enables reconstruction from distorted C-alpha-traces and multi-chain protein systems and features an optional fast minimization step which leads to further improvement of the reconstructed backbone atoms (other than C-alpha). Thanks to the included SCWRL4 library [42], the PD2 package also enables automatic reconstruction of side chain positions, so it is possible to get all-atomic structure from the alpha-carbon trace in one step. The PD2 method is available as a web-server and C++ standalone application at http://www.sbg.bio.ic.ac.uk/~phyre2/PD2_ca2main/.

In this work, we used the standalone version of the PD2 program to model backbone and C-beta atoms with additional minimization (*--ca2main:bb_min_steps* flag). The automatically reconstructed side chain atoms were rejected (to be automatically rebuilt again in early STAGE 3 of FlexPepDock refinement according to the Rosetta energy function). The command line used to rebuild atomic details from the C-alpha-trace of a single model is provided below:

~~~
~/bin/pd2_ca2main --database *./database/* -i *input.pdb -o output.pdb*
--ca2main:new_fixed_ca --ca2main:bb_min_steps *500*
~~~

The backbone or all-atom reconstruction stage of the modeling process can be also performed using other tools for protein structure reconstruction from the C-alpha trace [43]. It should be emphasized that proper reconstruction of structural details is crucial for effective model refinement and scoring reliability. Importantly, the tool used in STAGE 3 of the protocol (Rosetta FlexPepDock), can be very sensitive to model quality and local errors in the input structures.

### STAGE 3: Structure refinement using Rosetta FlexPepDock

Rosetta FlexPepDock is a well benchmarked and widely used tool for the high-resolution refinement of peptide-protein complexes [23, 44]. As demonstrated in several studies, it enables obtaining sub-Angstrom quality of protein-peptide structures [22, 44, 45]. The initial structure may be provided as a coarse-grained (at least backbone) or a fine-grained model, where the bound peptide position related to the receptor is relatively correct. The refinement accuracy depends not only on the spatial protein-peptide relationship but also on the initial peptide backbone conformation (e.g. its secondary structure). During refinement, FlexPepDock allows for full flexibility of the peptide while the receptor backbone is usually fixed and its side chains are optimized iteratively on-the-fly. The FlexPepDock tool is available as both an easy-to-use web-server (at http://flexpepdock.furmanlab.cs.huji.ac.il/) and a standalone protocol within the Rosetta package, which is freely available to academic users. The documentation and useful tutorials are provided at https://www.rosettacommons.org/.

FlexPepDock refinement consists of several operational modes that may be freely combined according to a specific task or input quality. The typical execution steps used for protein-peptide systems involve:

- preparation of properly formatted structure files,
- side chains pre-packing of initial complex structural components (removing internal clashes),
- refinement and/or minimization of pre-packed initial complex structure (optimization of protein-peptide interface interactions),
- selecting and clustering low-energy decoys (model scoring).

The details and technical hints for particular refinement steps are provided in the Supplementary ‘Technical details of the proposed workflow’ section.

In STAGE 3, we used the sets of 100 top-scored CABS-dock models reconstructed using PD2 (main chain and C-beta atoms only) as input for the FlexPepDock refinement procedure. Due to minor inaccuracies of local structure in coarse-grained models, which may have an adverse effect on overall Rosetta energy scoring, we reconstructed side chains and improved rotamer packing (*-flexpep_prepack* mode) directly at the Rosetta stage before taking further actions. Note that the FlexPepDock prepacking mode reconstructs automatically all missing side chain atoms in the initial receptor-peptide complex.

In the input sets of the top 100 models (resulting from STAGE 2), the average C-alpha RMSD for peptide molecules was in the range from 2.3 Å to 3.4 Å for six systems and from 3.1 Å to almost 8 Å for 3OE0 (based on comparison to its crystal structures). Therefore the sets included bound peptides, whose conformation and orientation in respect to the receptor was approximately correct. Using FlexPepDock, we tested two separate approaches to refine the sets of 100 top-scored models:

- STAGE 3: full refinement of the protein-peptide initial complex executed iteratively by Monte-Carlo search with energy minimization using the *-pep_refine* mode of the FlexPepDock protocol with optional fast low-resolution pre-optimization performed on coarse-grained *centroid* representation (average calculation time on a single CPU core: from 200 seconds to 350 seconds per model),
- STAGE 3’: short and fast minimization of the peptide-protein interface in the initial structure of the complex using the *-flexPepDockingMinimizeOnly* mode of the FlexPepDock protocol (average calculation time on a single CPU core: from 30 seconds to 40 seconds per model).

The command line used to reconstruct missing side chain atoms and pre-pack rotamers in a single initial peptide-protein complex (initial step of STAGE 3 and STAGE 3’) is as follows:

~~~
~FlexPepDocking.linuxgccrelease -database ${rosetta_db} -s *initial_prepack.pdb*
-native *native.pdb* -nstruct *1* -**flexpep_prepack** -ex1 -ex2aro -use_input_sc
~~~

The command line used to refine (main step of STAGE 3) the structure of a single initial peptide-protein complex is as follows:

~~~
~FlexPepDocking.linuxgccrelease -database ${rosetta_db} -s *initial_prepack.pdb*
-native *native.pdb* -out:file:silent *decoys.silent* -out:file:silent_struct_type *binary*
-nstruct *300* -**pep_refine** [-ex1 -ex2aro -use_input_sc -unboundrot file.pdb]^a^
[-detect_disulf *true* -rebuild_disulf *true* -fix_disulf *disulf.dat*]^b^ [-lowres_preoptimize]^c^
[-score:weights *talaris2014* -corrections::restore_talaris_behavior *true*]^d^
~~~

^a^ used in case the rotamer library is extended with extra side chain conformations

^b^ used in case the complex structure contains cysteines that may be connected by disulphide bridges

^c^ used in case of low-resolution pre-optimization

^d^ used in case of using the *talaris14* score function, default *REF2015*

The command line used to minimize the energy of a single initial peptide-protein complex (main step of STAGE 3’) is as follows:

~~~
~FlexPepDocking.linuxgccrelease -database ${rosetta_db} -s *initial_prepack.pdb*
-native *native.pdb* -out:file:silent *decoys.silent* -out:file:silent_struct_type *binary*-nstruct *1*
-**flexPepDockingMinimizeOnly** -ex1 -ex2aro -use_input_sc -detect_disulf *true*
-rebuild_disulf *true* -fix_disulf *disulf.dat*
~~~

The default output of FlexPepDock refinement (called by the *-out:file:silent* flag) is a silent file that contains the number of choice (called by the *-nstruct* flag) of compressed decoys and corresponding score values. The real Cartesian coordinates of selected decoys’ tags may be extracted by the built-in Rosetta protocol (*extract_pdbs*). The command line used for extraction of Cartesian coordinates is as follows:

~~~
~extract_pdbs.linuxgccrelease -database ${rosetta_db} -in:file:silent *file.silent*
-in:file:tags *file.txt*
~~~

Finally, the set of 10 top-scored models from STAGE 3 were selected using two-step hierarchical selection (initial selection of 1% top scored decoys form each refinement cycle according to *score* and *reweighted_sc* terms resulting in a set of 300 models, followed by selection of final 10 top models using *reweighted_sc* and *pep_sc* scoring terms). The set of 10 top-scored models from the alternative STAGE 3’ were selected from the set of 100 minimized model structures using *reweighted_sc* and *pep_sc* scoring terms. Technical details of the scoring procedure are provided in the Supplementary ‘Technical details of the proposed workflow’ section. The quality of the modeled receptor-peptide complexes was assessed by comparing their structural properties to corresponding crystal structures.

In the results analysis, we used the state-of-the-art measures proposed by the CAPRI assessors [46] to quantify various aspects of the quality of the models generated in the proposed protein-peptide docking protocol: IRMS (backbone root mean square deviation of the interface residues, after the interface components of the target and model have been superimposed), LRMS (backbone root mean square deviation of a peptide in the model relative to the target, after the receptor components of the target and model have been superimposed) and Fnat (the fraction of native residue–residue contacts).

All the technical information provided above is also included in the wiki pages of the CABS-dock standalone application under the following link: https://bitbucket.org/lcbio/cabsdock/wiki/Home#markdown-header-311-docking-to-gpcrs. This online information will be maintained and updated according to user’s needs.

## Results and discussion

In this work, we propose a novel and efficient protocol for the prediction of GPCR-peptide complex structures. The method was tested on a set of 7 GPCR-peptide complexes (see section GPCR benchmark dataset). In STAGE 1 (consisting of 10 independent CABS-dock simulations) the 100 top-scored coarse-grained models were selected out of a large set of 100,000 low energy models using structural clustering. The set of 100 top-scored models included receptor-peptide structures in C-alpha representation with medium and acceptable accuracy for 6 predicted complexes (based on ranking criteria for protein-peptide systems [46]).

For the 3OE0 system the best obtained model was ranked as incorrect showing a high IRMS value of 3.16 Å. During STAGE 2, the 100 top-scored models were reconstructed to main chain and C-beta atoms representation using the PD2 method [21]. In the final STAGE 3, models were subjected to the high-resolution FlexPepDock refinement procedure [22, 23] resulting in a set of 30,000 all-atom models for each predicted complex. At this point, we observed significant improvement in the models’ quality (see Table 1). The generated sets of models included medium and acceptable quality structures for 5 and 2 of modeled complexes, respectively. To deliver the best solution (i.e. a peptide-bound receptor model presenting the lowest IRMS value), in the small sets of 10 top-scored models, we used the procedure employing Rosetta scoring function (*REF2015: total score, reweighted_sc*) and interface-dependent energy terms (*pep_sc, I_sc*) [47, 48] (technical details of the scoring procedure are described in the Supplementary ‘Technical details of the proposed workflow’ section). Among the best selected-models, three complex structures presented medium accuracy, and another three models presented acceptable accuracy according to the CAPRI criteria for protein-peptide docking [46]. The best obtained model for the 3OE0 difficult case was ranked as inaccurate presenting an IRMS value of 2.41 Å. As an alternative for the computationally demanding refinement procedure (STAGE 3), we also tested a more straightforward approach (STAGE 3’) based on short and fast FlexPepDock minimization and scoring of top 100 reconstructed models (resulting from STAGE 2). The best minimized receptor-peptide structure among the 10 top-scored models for the studied complexes showed comparable quality to those resulting from STAGE 3 (see Table 1). Nevertheless, it should be emphasized that the most accurate atomistic models for each predicted complex were generated during the FlexPepDock refinement procedure (STAGE 3). The IRMS values obtained at each modeling stage are presented in Table 1, while corresponding LRMS and Fnat values are presented in Supplementary Table S3. Final structures of predicted receptor-peptide complexes (i.e. models showing the lowest IRMS values in the top 10 model sets resulting from STAGE 3 and STAGE 3’ of the modeling protocol) superimposed on corresponding crystal structures are presented in Figure 2.

**Table 1.**
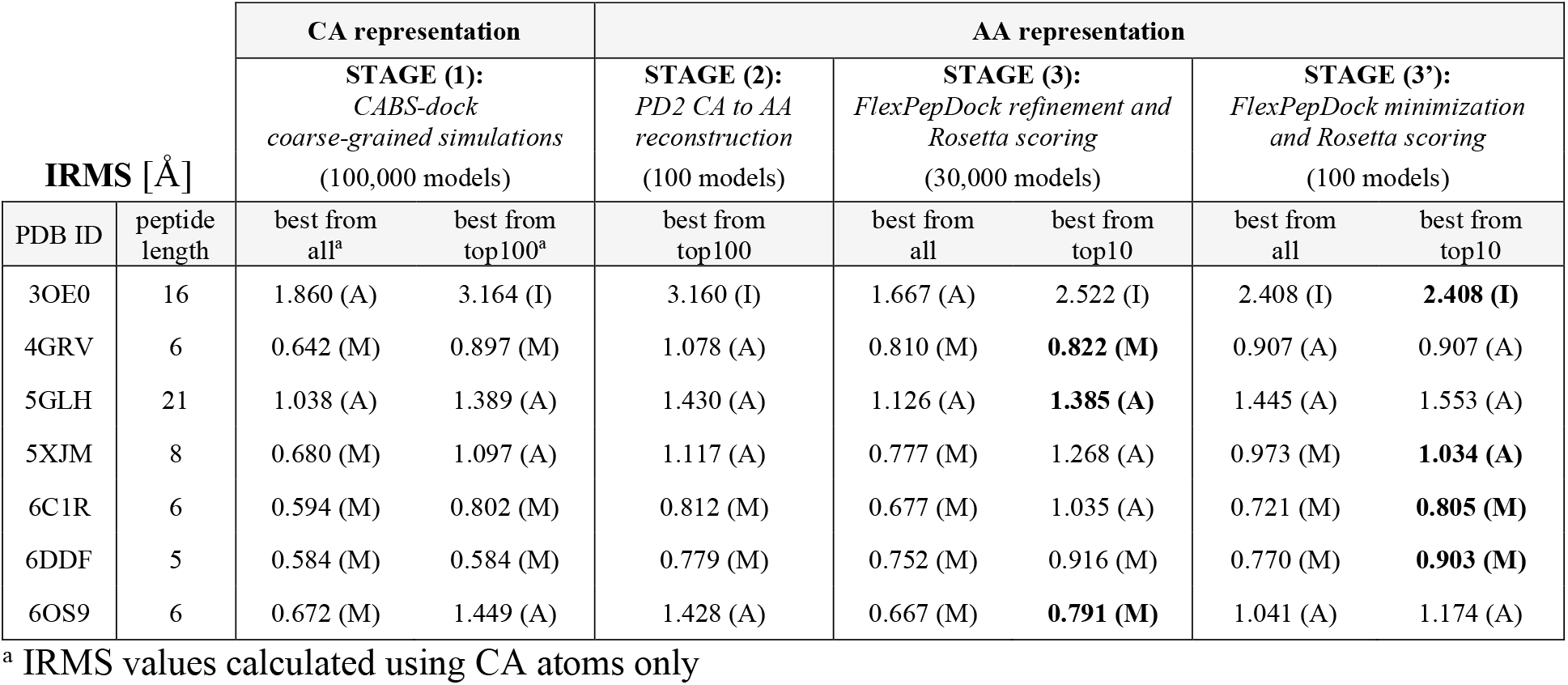
IRMS values calculated for docked peptides resulting from subsequent stages of the modeling procedure. Letters next to IRMS values indicate model accuracy and the following criteria were used: H, high (IRMS – 0 ≤ H ≤ 0.5 Å); M, medium (IRMS – 0.5 Å < M ≤ 1 Å); A, acceptable (IRMS – 1 Å < A ≤ 2 Å); I, incorrect (IRMS - 2 Å < I) (based on protein-peptide ranking criteria from the CAPRI experiment [46]).

**Figure 2.**
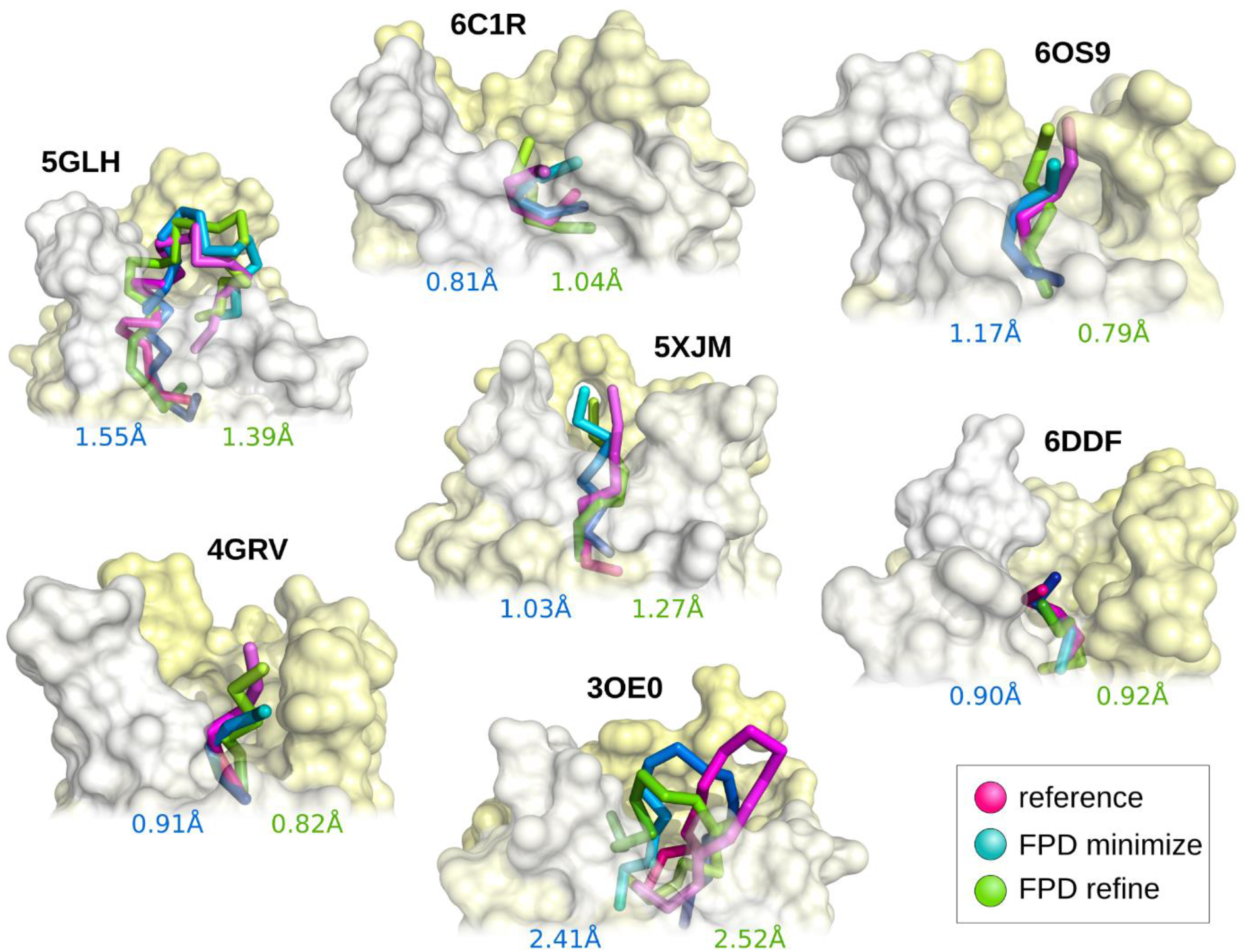
Comparison of the best predicted models (from the set of 10 top-scored models) with experimental peptide structures. Experimental models are presented in magenta, while predicted models in green (after refinement) and blue (after minimization).

The main goal of this work was to deliver an easy to use and efficient protocol for the accurate prediction of peptide ligand binding modes to the GPCR protein family. The protocol employs three state-of-the-art tools for the multiscale modeling of protein-peptide interactions [20–23, 42]. All these tools proved to be very effective for handling modeling tasks for many different systems consisting of cytosolic proteins. In this work, we propose their specific set-up dedicated for modeling GPCR-peptide systems. In STAGE 1 (docking of fully flexible peptides using CABS-dock) the peptide sampling space is restricted to spherical volume which includes all receptor fragments that may interact with bound peptides (for technical details see the Supplementary ‘Technical details of the proposed workflow’ section).

We also tested performance of the proposed protocol in which only single CABS-dock simulation was used for generation of initial top 10 models (STAGE 1 of the protocol) which were further subjected to the refinement and scoring procedure (see Supplementary Table S4). Results indicate that conducting only a single CABS-dock simulation run followed by FlexPepDock refinement also leads to good quality results, however slightly worse than that produced by combination of 10 independent simulation runs which enable much more efficient sampling of ligand binding poses.

It should be emphasized that the CABS-dock tool offers a simple way for using internal restraints in docked peptide molecules. This feature shows to be very useful especially for docking peptides containing internal disulfide bridges (e.g. endothelin, a 21-amino-acid peptide containing 2 disulfide bridges, docked to the ETB receptor, PDB ID: 5GLH). In addition, the application of restraints also proved to be very effective in preserving the cyclic conformation of the docked peptide (cyclic antagonist PMX53 docked to the C5a1 receptor, ODB ID: 6C1R).

One of the major challenges of the methods for protein-peptide docking is the scoring of a large set of models generated during predictions [6] because the score value does not always correlate with model quality. For optimal selection of the final 10 top-scored models resulting from STAGE 3 of the modeling protocol we used a two-step hierarchical approach. First we selected 1% of the best models according to *score* and *reweighted_sc* terms after each refine cycle: refinement of a single model resulting from STAGE 2 (where 1% accounted for the top 3 decoys selected from a set of 300 models generated during a single refinement cycle). This resulted in 300 pre-selected models for each modeled receptor-protein system. In the second step, final top 10 models were selected from 300 pre-selected decoys using *reweighted_sc* and *pep_sc* scores (for each term 5 top models were selected). The two-step scoring procedure described above provides maximized diversity of selected decoys and leads to much higher model accuracy when compared to selection of 1% of top scored models from the composite set of all 30000 decoys. The best of the obtained models in the sets of the 10 top-scored models are described using the IRMS values in Table 1, visualized in Figure 2 and characterized in Figure 3 (among the large pool of predicted models in terms of energy scores).

**Figure 3.**
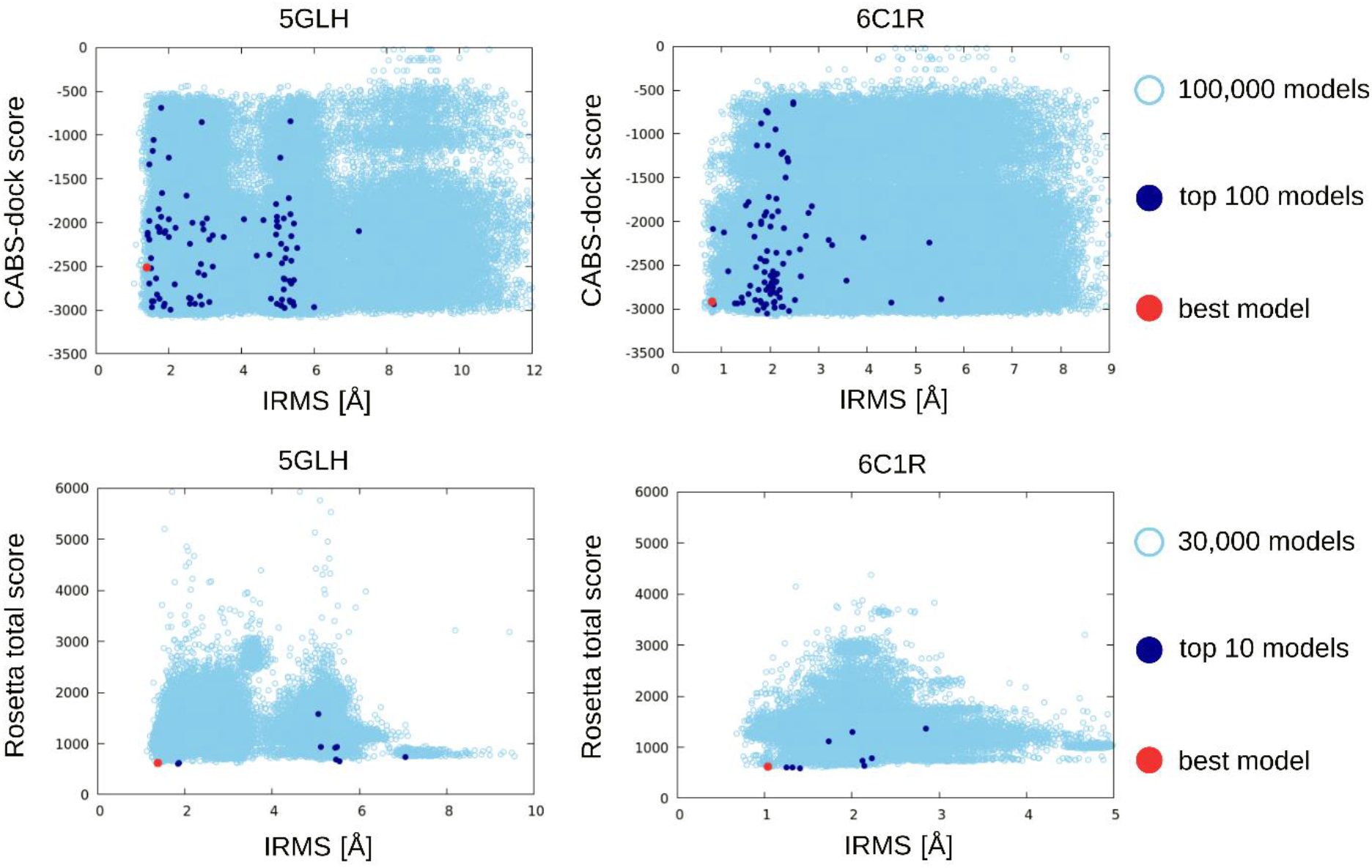
The plots show energy score versus IRMS [Å] for selected GPCR-peptide complexes (PDB ID: 5GLH, 6C1R). The first row contains CABS-dock results (STAGE 1, CA representation), while the second row contains final results after high-resolution FlexPepDock refinement (STAGE 3, AA representation). The top-scored models are colored in dark blue, and best models out of the top-scored models in red. The analogous plots for the other studied systems are provided in Supplementary Figure S1 and Figure S2.

## Conclusions

We have proposed and evaluated a novel three-stage protocol for prediction of GPCR-peptide complexes. The method employs CABS-dock docking simulations for generation of a large number of coarse-grained receptor-peptide complexes followed by PD2 model reconstruction to all-atom representation. Finally, Rosetta FlexPepDock is used for the model refinement and scoring procedure. The high-resolution solutions can be found in a small set of 10 top-scored models among the predicted structures. Application of the proposed protocol allowed for successful prediction of 6 out of 7 GPCR-peptide complexes providing structures with acceptable or medium accuracy (according to the criteria of CAPRI competition [46]). The proposed protocol can be easily extended to structure prediction exercises that support flexibility of selected GPCR binding fragments during docking simulations, which is possible in CABS-dock docking.

## Key points

- CABS-dock docking restricted to the broad neighborhood of the binding site enables flexible docking of peptides to membrane receptors with high accuracy.
- Rosetta FlexPepDock refinement of CABS-dock predicted GPCR-peptide complexes leads to significant improvement of model quality.
- We present and validate a protocol for CABS-dock docking and further FlexPepDock refinement and scoring allowing high-resolution prediction of GPCR-peptide complexes.

## Biographical notes

Aleksandra Badaczewska-Dawid is postdoc research associate at the Bioscience Innovation Postdoctoral Fellowship, Department of Chemistry, Iowa State University, Ames, USA.

Sebastian Kmiecik is Associate Professor at the Faculty of Chemistry, University of Warsaw, and leads a Laboratory of Computational Biology research group at the Biological and Chemical Research Centre, University of Warsaw, Poland.

Michał Koliński is the head of Bioinformatics Laboratory at the Mossakowski Medical Research Centre Polish Academy of Science, Warsaw, Poland.

## Acknowledgments

This work was supported by the National Science Centre of Poland, grant MAESTRO no. 2014/14/A/ST6/00088.

## References

1. Henninot A, Collins JC, Nuss JM, The Current State of Peptide Drug Discovery: Back to the Future? J Med Chem, 2018. 61(4): 1382–1414.

2. Muratspahić E, Freissmuth M, Gruber CW, Nature-derived peptides: a growing niche for GPCR ligand discovery. Trends in pharmacological sciences, 2019.

3. Wu F, Song G, de Graaf C, et al., Structure and Function of Peptide-Binding G Protein-Coupled Receptors. J Mol Biol, 2017. 429(17): 2726–2745.

4. Heydenreich FM, Vuckovic Z, Matkovic M, et al., Stabilization of G protein-coupled receptors by point mutations. Front Pharmacol, 2015. 6: 82.

5. Pándy-Szekeres G, Munk C, Tsonkov TM, et al., GPCRdb in 2018: adding GPCR structure models and ligands. Nucleic acids research, 2018. 46(D1): D440–D446.

6. Ciemny M, Kurcinski M, Kamel K, et al., Protein-peptide docking: opportunities and challenges. Drug Discov Today, 2018. 23(8): 1530–1537.

7. Sandal M, Duy TP, Cona M, et al., GOMoDo: A GPCRs online modeling and docking webserver. PLoS One, 2013. 8(9): e74092.

8. Launay G, Teletchea S, Wade F, et al., Automatic modeling of mammalian olfactory receptors and docking of odorants. Protein Eng Des Sel, 2012. 25(8): 377–86.

9. Lee GR, Seok C, Galaxy7TM: flexible GPCR–ligand docking by structure refinement. Nucleic acids research, 2016. 44(W1): W502–W506.

10. Bartuzi D, Kaczor AA, Targowska-Duda KM, et al., Recent Advances and Applications of Molecular Docking to G Protein-Coupled Receptors. Molecules, 2017. 22(2): 340.

11. Rentzsch R, Renard BY, Docking small peptides remains a great challenge: an assessment using AutoDock Vina. Briefings in Bioinformatics, 2015. 16(6): 1045–1056.

12. Wedemeyer MJ, Mueller BK, Bender BJ, et al., Comparative modeling and docking of chemokine-receptor interactions with Rosetta. Biochem Biophys Res Commun, 2020.

13. Zachmann J, Kritsi E, Tapeinou A, et al., Combined Computational and Structural Approach into Understanding the Role of Peptide Binding and Activation of Melanocortin Receptor 4. J Chem Inf Model, 2020.

14. Sencanski M, Glisic S, Snajder M, et al., Computational design and characterization of nanobody-derived peptides that stabilize the active conformation of the beta2-adrenergic receptor (beta2-AR). Sci Rep, 2019. 9(1): 16555.

15. Kufareva I, Handel TM, Abagyan R, Experiment-guided molecular modeling of protein–protein complexes involving GPCRs, in G Protein-Coupled Receptors in Drug Discovery. 2015, Springer. 295–311.

16. Karhu L, Turku A, Xhaard H, Modeling of the OX 1 R–orexin-A complex suggests two alternative binding modes. BMC structural biology, 2015. 15(1): 9.

17. Thibeault PE, LeSarge JC, Fernandes M, et al., Molecular basis for activation and biased signalling at the thrombin-activated GPCR Proteinase Activated Receptor-4 (PAR4). Journal of Biological Chemistry, 2019: jbc. RA119. 011461.

18. Peeters MC, van Westen GJ, Li Q, et al., Importance of the extracellular loops in G protein-coupled receptors for ligand recognition and receptor activation. Trends Pharmacol Sci, 2011. 32(1): 35–42.

19. Kolinski M, Plazinska A, Jozwiak K, Recent progress in understanding of structure, ligand interactions and the mechanism of activation of the β 2-adrenergic receptor. Current medicinal chemistry, 2012. 19(8): 1155–1163.

20. Kurcinski M, Pawel Ciemny M, Oleniecki T, et al., CABS-dock standalone: a toolbox for flexible protein-peptide docking. Bioinformatics, 2019. 35(20): 4170–4172.

21. Moore BL, Kelley LA, Barber J, et al., High–quality protein backbone reconstruction from alpha carbons using Gaussian mixture models. J Comput Chem, 2013. 34(22): 1881–1889.

22. Raveh B, London N, Schueler-Furman O, Sub-angstrom modeling of complexes between flexible peptides and globular proteins. Proteins, 2010. 78(9): 2029–40.

23. London N, Raveh B, Cohen E, et al., Rosetta FlexPepDock web server--high resolution modeling of peptide-protein interactions. Nucleic Acids Res, 2011. 39(Web Server issue): W249–53.

24. Tikhonova IG, Gigoux V, Fourmy D, Understanding peptide binding in Class AG protein-coupled receptors. Molecular pharmacology, 2019. 96(5): 550–561.

25. Wu B, Chien EY, Mol CD, et al., Structures of the CXCR4 chemokine GPCR with small-molecule and cyclic peptide antagonists. Science, 2010. 330(6007): 1066–71.

26. White JF, Noinaj N, Shibata Y, et al., Structure of the agonist-bound neurotensin receptor. Nature, 2012. 490(7421): 508–13.

27. Shihoya W, Nishizawa T, Okuta A, et al., Activation mechanism of endothelin ET B receptor by endothelin-1. Nature, 2016. 537(7620): 363–368.

28. Asada H, Horita S, Hirata K, et al., Crystal structure of the human angiotensin II type 2 receptor bound to an angiotensin II analog. Nat Struct Mol Biol, 2018. 25(7): 570–576.

29. Liu H, Kim HR, Deepak R, et al., Orthosteric and allosteric action of the C5a receptor antagonists. Nat Struct Mol Biol, 2018. 25(6): 472–481.

30. Koehl A, Hu H, Maeda S, et al., Structure of the μ-opioid receptor–G i protein complex. Nature, 2018. 558(7711): 547–552.

31. Kato HE, Zhang Y, Hu H, et al., Conformational transitions of a neurotensin receptor 1–G i1 complex. Nature, 2019. 572(7767): 80–85.

32. Kurcinski M, Badaczewska-Dawid A, Kolinski M, et al., Flexible docking of peptides to proteins using CABS-dock. Protein Sci, 2020. 29(1): 211–222.

33. Kmiecik S, Kouza M, Badaczewska-Dawid AE, et al., Modeling of Protein Structural Flexibility and Large-Scale Dynamics: Coarse-Grained Simulations and Elastic Network Models. Int J Mol Sci, 2018. 19(11): 3496.

34. Ciemny MP, Badaczewska-Dawid AE, Pikuzinska M, et al., Modeling of disordered protein structures using monte carlo simulations and knowledge-based statistical force fields. International journal of molecular sciences, 2019. 20(3): 606.

35. Kmiecik S, Gront D, Kolinski M, et al., Coarse-grained protein models and their applications. Chemical reviews, 2016. 116(14): 7898–7936.

36. Kurcinski M, Jamroz M, Blaszczyk M, et al., CABS-dock web server for the flexible docking of peptides to proteins without prior knowledge of the binding site. Nucleic Acids Res, 2015. 43(W1): W419–24.

37. Blaszczyk M, Kurcinski M, Kouza M, et al., Modeling of protein-peptide interactions using the CABS-dock web server for binding site search and flexible docking. Methods, 2016. 93: 72–83.

38. Ciemny MP, Kurcinski M, Blaszczyk M, et al., Modeling EphB4-EphrinB2 protein-protein interaction using flexible docking of a short linear motif. Biomed Eng Online, 2017. 16(Suppl 1): 71.

39. Ciemny MP, Debinski A, Paczkowska M, et al., Protein-peptide molecular docking with large-scale conformational changes: the p53-MDM2 interaction. Sci Rep, 2016. 6: 37532.

40. Blaszczyk M, Ciemny MP, Kolinski A, et al., Protein-peptide docking using CABS-dock and contact information. Brief Bioinform, 2019. 20(6): 2299–2305.

41. Koliński M, Kmiecik S, Dec R, et al., Docking interactions determine early cleavage events in insulin proteolysis by pepsin: Experiment and simulation. International Journal of Biological Macromolecules, 2020.

42. Krivov GG, Shapovalov MV, Dunbrack RL, Jr., Improved prediction of protein side-chain conformations with SCWRL4. Proteins, 2009. 77(4): 778–95.

43. Badaczewska-Dawid AE, Kolinski A, Kmiecik S, Computational reconstruction of atomistic protein structures from coarse-grained models. Comput Struct Biotechnol J, 2020. 18: 162–176.

44. Alam N, Schueler-Furman O, Modeling Peptide-Protein Structure and Binding Using Monte Carlo Sampling Approaches: Rosetta FlexPepDock and FlexPepBind. Methods Mol Biol, 2017. 1561: 139–169.

45. Liu T, Pan X, Chao L, et al., Subangstrom accuracy in pHLA-I modeling by Rosetta FlexPepDock refinement protocol. J Chem Inf Model, 2014. 54(8): 2233–42.

46. Lensink MF, Velankar S, Wodak SJ, Modeling protein–protein and protein–peptide complexes: CAPRI 6th edition. Proteins: Structure, Function, and Bioinformatics, 2017. 85(3): 359–377.

47. Leaver-Fay A, O’Meara MJ, Tyka M, et al., Scientific benchmarks for guiding macromolecular energy function improvement, in Methods in enzymology. 2013, Elsevier. 109–143.

48. Alford RF, Leaver-Fay A, Jeliazkov JR, et al., The Rosetta All-Atom Energy Function for Macromolecular Modeling and Design. J Chem Theory Comput, 2017. 13(6): 3031–3048.

