## Supplementary information for "Docking of peptides to GPCRs using a combination of CABS-dock with FlexPepDock refinement"

### Contents:

Technical details of the proposed workflow

Table S1

Table S2

Table S3

Table S4

Figure S1

Figure S2

SI References

### Technical details of the proposed workflow

STAGE 1: CABS-dock coarse-grained simulations

*Definition of the docking sphere*

The distance restraints, limiting the docking area to the broad neighborhood of the binding site and N-terminal domain of the receptor, were defined independently for each system. The following procedure was applied. First, the receptor residue positioned near the center of the binding site was selected. Next, restraints were imposed between the side chain (SC) atom of the selected receptor residue and each SC atom of the docked ligand. The restraint distances were adjusted to the size of the docked peptide, namely two arbitrary values were selected: 25 Å for shorter peptides (below 7 residues) and 30 Å for two the longest peptides (above 15 residues), respectively. In this way the peptide sampling volume was reduced to the sphere (with a radius from 25 Å to 30 Å) centered on the receptor binding site. The distance restraints are introduced into the CABS-energy function as an additional contact energy term, given by the formula:

$$E_{contact}(d) = \begin{cases} 0 \; if \; d \leq D_0 \\ s(d - D_0) \; if \; d > D_0 \end{cases}$$

where $d$ is the distance between the centers of mass of two restrained side-chains (SC), $D_0$ is the distance cut-off and $s$ is the weight of the restraint. If the distance between the restrained residues exceeds the user-defined threshold $D_0$ (25 Å or 30 Å in our case), the energetic penalty linearly increases with the slope defined by the restraint weight, $s$ (default: 5.0). If the measured distance is below the cut-off, peptide scoring is not affected. The detailed information used for docking sphere parameters is given in Table S1. The command lines used in this study for CABS-dock docking simulations are presented in Table S2.

STAGE 2: PD2 structure reconstruction to all-atom representation

After successful compilation with SCons (software construction tool, see https://scons.org/), the PD2 executables (e.g. *pd2_ca2main*) are available in the *bin/* directory. When further backbone minimization is needed, the number of cycles for the option *--ca2main:bb_min_steps* has to be determined (500 is the recommended value). The procedure is fully automatic and generates all-atom reconstructed structure as well as a short summary on the standard output stream. Everything is provided: score line with contributions of various force field components, elapsed time and validation of output structure.

STAGE 3: Rosetta FlexPepDock high-resolution refinement

*Input preparation*

Input preparation is an important step that can strongly affect the success and accuracy of the refinement. The obligatory input to the refinement protocol is an initial peptide-receptor complex structure provided in a single file of the PDB format, either with or without side-chain coordinates (backbone atoms only are sufficient). There are several simple rules that must be fulfilled: the receptor is always the first chain in the file, no heteroatoms or double conformations are included, the occupancy column is filled (no 0.00 values are allowed), the numbering of residues in the receptor and peptide is continuous and starting from 1 (disregards PDB numbering). The same rules apply to the numbering of constraints provided in separate files. The recommended way to get a correctly formatted starting complex structure is to use a ready-made Rosetta python script *clean_pdb.py* that can be found on the path tools/protein_tools/scripts/clean_pdb.py in the local Rosetta copy. Additional inputs that may be provided optionally include: known complex structure (e.g. experimental or constructed from a homology model) as a reference for RMSD calculations; unbound receptor structure that increases the side-chain rotamer library and atom indices pair type constraints files that define e.g. contacts or disulphide bridges. The FlexPepDock protocol has been developed to improve the starting peptide conformation (even the extended peptide backbone) that is positioned approximately close to the correct binding site (despite the wrong spatial orientation). It is highly recommended to discard the input side-chains from bound-type native complexes, but to keep the side-chains from unbound receptor structures and provide additional rotamer flags (-ex1 and -ex2aro), which increases the rotamer library and may improve final bound structures.

*Prepacking the side chains of the initial complex*

The side chains in both peptide and receptor initial structures can contain internal clashes, especially if models were obtained using lower-resolution coarse-grained tools. Those inaccuracies of the local structure may have an adverse effect on overall Rosetta energy scoring, which is used for ranking decoys. This applies not only to the binding interface but also to the entire protein-peptide complex structure. Therefore, it is highly recommended to improve rotamer packing before the refinement stage. For this purpose, the Rosetta package offers a dedicated *-flexpep_prepack* mode that optimizes the side-chains of each residue according to the chosen Rosetta scoring function (default *REF2015*). The suggested rotamers can be determined by the *-use_input_sc* flag in case where side-chains are available in the input structure (e.g. reconstructed or derived from homology modeling) and/or by -the *unboundrot* flag when the unbound structure is known.

*Refinement of the pre-packed protein-peptide complex*

The main step of the FlexPepDock protocol is a refinement mode called by the *-pep_refine* flag. During that stage the pre-packed initial complex structure is iteratively refined by Monte-Carlo search with energy minimization. In particular, the receptor backbone out of the binding site remains almost fixed, while the rigid-body orientation of the peptide relative to the receptor and its backbone undergo perturbations. The interface side-chains rotamers of both the peptide and the receptor have full flexibility and are periodically optimized, while in the final cycle the minimization procedure is applied to all side chains in the system. The attractive and repulsive components of van der Waals terms are gradually scaled (toward their standard values) both to ensure efficient perturbations at the binding site and to avoid peptide-protein spatial separation. The FlexPepDock protocol also provides an optional step of fast low-resolution pre-optimization performed on coarse-grained *centroid* representation (the *lowres_preoptimize* flag) and minimization according to smoothened *score4* energy function. This may additionally improve the conformation of the initial complex before high-resolution refinement, especially when the initial complex structure shows backbone-RMSD values larger than 3 Å when compared to the reference structure.

Based on the experience of the FlexPepDock developers, we performed two separate runs with and without the -*lowres_preoptimize* flag, and each generated 300 refined decoys. Due to the lack of known unbound receptor structures for all studied systems, only the suggested extra rotamer flags (*-ex1 and -ex2aro*) were used. Since most of the protein-peptide complexes studied in this work contained pairs of cysteines bounded by a disulphide bridge, we used additional options for the automatic detection of and rebuilding disulphide connections (-*etect_disulf true* and *-rebuild_disulf true*). In addition, for the two systems (3OE0, 5GLH) disulphide bridges in docked peptides were defined as constraints (*-fix_disulf* flag) provided in a separate file (*disulf.dat*) as a pair of residue indices per single line. We tested protocols with both the *talaris2014* and *REF2015* energy score, while the results from the second series of simulations are presented only, due to their much higher accuracy.

*Scoring procedure after refinement and minimization*

The selection of the final models among all the refined decoys is usually a challenging step due to the unavailability of ideal correlation between the scoring function and model quality. The composite set of all the decoys obtained from FlexPepDock refinement is usually ranked according to both the default *total score* and/or *reweighted-score* (where the weights of interface residues and peptide residues are increased) and/or clustered using Rosetta cluster application (*cluster.py*).

Since the set of 10 top-scored models, chosen by *total score* or *reweighted score* from (a) 30000 refined decoys or (b) representative models selected from low-energy clusters (of 1-2 Å in radius) was generally of lower quality than prepacked initial complex structures, we performed an additional analysis of score terms related to internal peptide energy (*pep_sc*) and interface energy (*I_sc*) for their ability to identify the near-native models. The highest ranking success (for all studied receptor-peptide complexes) was achieved by two-step hierarchical selection, which was used for the selection of the final top 10 models of STAGE 3, where:

- In the first step, we selected 1% top-scored models (according to *score* and *reweighted_sc* terms) obtained independently from each refine cycle (where 1% accounted for top 3 decoys selected from a set of 300 models generated during a single refine cycle conducted on a single receptor-peptide model). Since the set of the 100 top-scored models (resulting from STAGE 2) for each complex were submitted to the refined procedure, top 3 × 100 = 300 models were selected. This step enables maximized diversity of selected models and provided much higher model accuracy when compared to selection of 1% of top scored models from the composite set of all 30000 decoys.
- In the second step we ranked the pre-selected 300 models (selected during the first step described above) according to *reweighted_sc* and *pep_sc* terms. For each term 5 top models were selected resulting in a composite set of top 10 final STAGE 3 models.

The set of 10 top-scored models resulting from STAGE 3’ were selected using the second step of the procedure described above only. That is, the sets of 100 minimized models were ranked according to *reweighted_sc* and *pep_sc* terms and top 5 + 5 = 10 models were selected.

**Table S1.** Structural data including peptide sequence information and restraints parameters used for CABS-dock input during test predictions of GPCR-peptide complexes.

| PDB ID | peptide length | docked peptide sequence | substituted residue(s) in docked peptides | restrained residues of peptide* | docking sphere parameters: |
|---|---|---|---|---|---|
| code | int | string | (X) | [pep-res/distance, Å] | [rec-res/distance, Å] |
| 5GLH | 21 | CSCSSLMDKECV YFCHLDIIW | - | Cys1-Cys15/5.3A; Cys3-Cys11/6.2A | 249:A/30A |
| 4GRV | 6 | RRPYIL | - | - | 215:A/25A |
| 5XJM | 8 | GRVYIHPI | (X)RVYIHPI | - | 155:A/25A |
| 6OS9 | 6 | KKPYIL | - | - | 163:A/25A |
| 6DDF | 5 | YAGFG | Y(X)G(X)(X) | - | 147:R/25A |
| 6C1R | 6 | FAPLWR | F(X)P(X)WR | Ala2-Arg6/6.09A (cyclic peptide) | 153:B/25A |
| 3OE0 | 16 | RRWCYQKPPYRR CRGP | RR(X)CYQK(X)P YRRCRG(X) | Cys4-Cys13/3.9A | 3:A/30A |

*the restraint distances were derived from crystal structures PDB ID: 6DK5 (for endothelin-1 peptide docked to the $ET_B$ receptor and PDB ID: 3OE0 (for antagonist CVX15 docked to the CXCR4 receptor). In the case of 6C1R structure the distance restrains were imposed between CA atoms of Arg2 and Arg6 of the docked peptide to preserve its cyclic conformation.

**Table S2.** CABS-dock input command lines and associated structure files used for prediction of GPCR-peptide complexes. Structure files were created based on corresponding crystal structures derived from the PDB database. Data enables reproduction of CABS-dock docking results presented in this study. A detailed description of input CABS-dock commands presented below is available at https://bitbucket.org/lcbio/cabsdock/wiki/.

| | |
|---|---|
| **3OE0** | Structure file: 3OE0_struc.pdb (receptor - chain A; peptide ligand - chain I) |
| | *CABS-dock input command:*<br>CABSdock -s 100 -M -C -S -v 4 -i 3OE0_struc.pdb:A -p 3OE0_struc.pdb:I --reference-pdb 3OE0_struc.pdb:A:I --ca-rest-add 4:PEP 13:PEP 3.9 5.0 --sc-rest-add 3:A 1:PEP 30.0 5.0 --sc-rest-add 3:A 2:PEP 30.0 5.0 --sc-rest-add 3:A 3:PEP 30.0 5.0 --sc-rest-add 3:A 4:PEP 30.0 5.0 --sc-rest-add 3:A 5:PEP 30.0 5.0 --sc-rest-add 3:A 6:PEP 30.0 5.0 --sc-rest-add 3:A 7:PEP 30.0 5.0 --sc-rest-add 3:A 8:PEP 30.0 5.0 --sc-rest-add 3:A 9:PEP 30.0 5.0 --sc-rest-add 3:A 10:PEP 30.0 5.0 --sc-rest-add 3:A 11:PEP 30.0 5.0 --sc-rest-add 3:A 12:PEP 30.0 5.0 --sc-rest-add 3:A 13:PEP 30.0 5.0 --sc-rest-add 3:A 14:PEP 30.0 5.0 --sc-rest-add 3:A 15:PEP 30.0 5.0 --sc-rest-add 3:A 16:PEP 30.0 5.0 |
| **4GRV** | Structure file: 4GRV_struc.pdb (receptor - chain A ; peptide ligand - chain B ) |
| | *CABS-dock input command:*<br>CABSdock -s 100 -M -C -S -v 4 -i 4GRV_struc.pdb:A -p 4GRV_struc.pdb:B --reference-pdb 4GRV_struc.pdb:A:B --sc-rest-add 215:A 1:PEP 25.0 5.0 --sc-rest-add 215:A 2:PEP 25.0 5.0 --sc-rest-add 215:A 3:PEP 25.0 5.0 --sc-rest-add 215:A 4:PEP 25.0 5.0 --sc-rest-add 215:A 5:PEP 25.0 5.0 --sc-rest-add 215:A 6:PEP 25.0 5.0 |
| **5GLH** | Structure file: 5GLH_struc.pdb (receptor - chain A; peptide ligand - chain B) |
| | *CABS-dock input command:*<br>CABSdock -s 100 -M -C -S -v 4 -i 5GLH_struc.pdb:A -p 5GLH_struc.pdb:B --reference-pdb 5GLH_struc.pdb:A:B --ca-rest-add 1:PEP 15:PEP 5.3 1.0 --ca-rest-add 3:PEP 11:PEP 6.2 1.0 --sc-rest-add 249:A 1:PEP 30.0 5.0 --sc-rest-add 249:A 2:PEP 30.0 5.0 --sc-rest-add 249:A 3:PEP 30.0 5.0 --sc-rest-add 249:A 4:PEP 30.0 5.0 --sc-rest-add 249:A 5:PEP 30.0 5.0 --sc-rest-add 249:A 6:PEP 30.0 5.0 --sc-rest-add 249:A 7:PEP 30.0 5.0 --sc-rest-add 249:A 8:PEP 30.0 5.0 --sc-rest-add 249:A 9:PEP 30.0 5.0 --sc-rest-add 249:A 10:PEP 30.0 5.0 --sc-rest-add 249:A 11:PEP 30.0 5.0 --sc-rest-add 249:A 12:PEP 30.0 5.0 --sc-rest-add 249:A 13:PEP 30.0 5.0 --sc-rest-add 249:A 14:PEP 30.0 5.0 --sc-rest-add 249:A 15:PEP 30.0 5.0 --sc-rest-add 249:A 16:PEP 30.0 5.0 --sc-rest-add 249:A 17:PEP 30.0 5.0 --sc-rest-add 249:A 18:PEP 30.0 5.0 --sc-rest-add 249:A 19:PEP 30.0 5.0 --sc-rest-add 249:A 20:PEP 30.0 5.0 --sc-rest-add 249:A 21:PEP 30.0 5.0 |
| **5XJM** | Structure file: 5XJM_struc.pdb (receptor - chain A; peptide ligand - chain B ) |
| | *CABS-dock input command:*<br>CABSdock -s 100 -M -C -S -v 4 -i 5XJM_struc.pdb:A -p 5XJM_struc.pdb:B --reference-pdb 5XJM_struc.pdb:A:B --sc-rest-add 155:A 1:PEP 25.0 5.0 --sc-rest-add 155:A 2:PEP 25.0 5.0 --sc-rest-add 155:A 3:PEP 25.0 5.0 --sc-rest-add 155:A 4:PEP 25.0 5.0 --sc-rest-add 155:A 5:PEP 25.0 5.0 --sc-rest-add 155:A 6:PEP 25.0 5.0 --sc-rest-add 155:A 7:PEP 25.0 5.0 --sc-rest-add 155:A 8:PEP 25.0 5.0 |
| **6C1R** | Structure file: 6C1R_struc.pdb (receptor - chain B; peptide ligand - chain L) |
| | *CABS-dock input command:*<br>CABSdock -s 100 -M -C -S -v 4 -i 6C1R_struc.pdb:B -p 6C1R_struc.pdb:L --reference-pdb 6C1R_struc.pdb:B:L --ca-rest-add 2:PEP 6:PEP 6.09 5.0 --sc-rest-add 153:B 1:PEP 25.0 5.0 --sc-rest-add 153:B 2:PEP 25.0 5.0 --sc-rest-add 153:B 3:PEP 25.0 5.0 --sc-rest-add 153:B 4:PEP 25.0 5.0 --sc-rest-add 153:B 5:PEP 25.0 5.0 --sc-rest-add 153:B 6:PEP 25.0 5.0 |
| **6DDF** | Structure file: 6DDF_struc.pdb (receptor - chain R; peptide ligand - chain D) |
| | *CABS-dock input command:*<br>CABSdock -s100 -M -C -S -v 4 -i 6DDF_struc.pdb:R -p 6DDF_struc.pdb:D --reference-pdb 6DDF_struc.pdb:R:D --sc-rest-add 147:R 1:PEP 25.0 5.0 --sc-rest-add 147:R 2:PEP 25.0 5.0 --sc-rest-add 147:R 3:PEP 25.0 5.0 --sc-rest-add 147:R 4:PEP 25.0 5.0 --sc-rest-add 147:R 5:PEP 25.0 5.0 |
| **6OS9** | Structure file: 6OS9_struc.pdb (receptor - chain R; peptide ligand - chain L) |
| | *CABS-dock input command:*<br>CABSdock -s 100 -M -C -S -v 4 -i 6OS9_struc.pdb:R -p 6OS9_struc.pdb:L --reference-pdb 6OS9_struc.pdb:R:L --sc-rest-add 163:R 1:PEP 25.0 5.0 --sc-rest-add 163:R 2:PEP 25.0 5.0 --sc-rest-add 163:R 3:PEP 25.0 5.0 --sc-rest-add 163:R 4:PEP 25.0 5.0 --sc-rest-add 163:R 5:PEP 25.0 5.0 --sc-rest-add 163:R 6:PEP 25.0 5.0 |

**Table S3.** LRMS and Fnat values calculated for predicted models of GPCR-peptide complexes using the proposed modeling protocol. Data correspond to IRMS values given in Table 1 in the manuscript. All the values were calculated using protein-peptide ranking criteria from the CAPRI experiment [1].

| | | **CA representation** | | **AA representation** | | | | |
|---|---|---|---|---|---|---|---|---|
| | | **STAGE (1):** *CABS-dock coarse-grained simulations* (100,000 models) | | **STAGE (2):** *PD2 CA to AA reconstruction* (100 models) | **STAGE (3):** *FlexPepDock refinement and Rosetta scoring* (30,000 models) | | **STAGE (3'):** *FlexPepDock minimization and Rosetta scoring* (100 models) | |
| **LRMS** [Å] | | | | | | | | |
| PDB ID | peptide length | best from all[a] | top100[a] | top100 | best from all | top10 | best from all | top10 |
| 3OE0 | 16 | 5.117 | 9.103 | 9.075 | 4.413 | 6.899 | 6.660 | 6.660 |
| 4GRV | 6 | 1.932 | 3.130 | 3.605 | 2.041 | 1.987 | 2.726 | 2.726 |
| 5GLH | 21 | 2.627 | 3.576 | 3.571 | 2.633 | 3.604 | 3.650 | 4.010 |
| 5XJM | 8 | 2.036 | 4.058 | 4.031 | 2.257 | 4.643 | 3.250 | 3.555 |
| 6C1R | 6 | 1.896 | 2.602 | 2.532 | 2.015 | 3.650 | 2.190 | 2.479 |
| 6DDF | 5 | 1.487 | 1.487 | 2.215 | 1.954 | 2.891 | 2.139 | 3.153 |
| 6OS9 | 6 | 2.034 | 5.616 | 5.536 | 1.381 | 2.266 | 3.302 | 4.285 |
| **Fnat** [Å] | | best from all[a] | top100[a] | top100 | best from all | top10 | best from all | top10 |
| 3OE0 | 16 | 0.017 | 0.103 | 0.103 | 0.293 | 0.310 | 0.293 | 0.293 |
| 4GRV | 6 | 0.018 | 0.127 | 0.127 | 0.509 | 0.527 | 0.582 | 0.582 |
| 5GLH | 21 | 0.009 | 0.102 | 0.074 | 0.398 | 0.343 | 0.361 | 0.315 |
| 5XJM | 8 | 0.000 | 0.078 | 0.078 | 0.569 | 0.275 | 0.431 | 0.373 |
| 6C1R | 6 | 0.021 | 0.271 | 0.292 | 0.562 | 0.479 | 0.604 | 0.542 |
| 6DDF | 5 | 0.040 | 0.120 | 0.120 | 0.680 | 0.440 | 0.640 | 0.480 |
| 6OS9 | 6 | 0.000 | 0.120 | 0.120 | 0.580 | 0.500 | 0.460 | 0.360 |

[a] IRMS values calculated using CA atoms only

**Table S4**. The lowest IRMS values (and corresponding LRMS and Fnat values) obtained for top 10 GPCR-peptide complex models predicted using the modified three-stage modeling protocol (using only single CABS-dock simulation during STAGE 1). The subsequent modeling tasks included: STAGE 1, single CABS-dock simulation resulting in top 10 models; STAGE 2, reconstruction of top 10 models using PD2; STAGE 3, full refinement of protein-peptide complex models executed iteratively by Monte-Carlo search with energy minimization (resulting in a 3000 model set) and scoring (resulting in top 10 final models). Results are presented for 10 independent protocol executions (P1 to P10) for each of 7 GPCR-peptide complexes analyzed in this study.

| Lowest IRMS values for top 10 models (10/3000) for 10 independent protocol executions [Å] | | | | | | | | | | |
|---|---|---|---|---|---|---|---|---|---|---|
| PDB ID | P1 | P2 | P3 | P4 | P5 | P6 | P7 | P8 | P9 | P10 |
| **3OE0** | 3.659 | 2.522 | 3.668 | 3.953 | 3.078 | 3.300 | 3.594 | 2.900 | 3.484 | 2.698 |
| **4GRV** | 1.147 | 0.965 | 1.233 | 0.822 | 1.102 | 0.936 | 1.078 | 0.965 | 1.389 | 0.972 |
| **5GLH** | 1.596 | 1.508 | 2.343 | 1.613 | 1.385 | 1.508 | 5.226 | 1.641 | 1.236 | 2.069 |
| **5XJM** | 1.060 | 1.272 | 1.058 | 1.381 | 1.268 | 1.165 | 1.367 | 1.153 | 1.254 | 1.172 |
| **6C1R** | 1.388 | 1.723 | 1.560 | 1.035 | 1.833 | 1.925 | 1.880 | 1.659 | 2.057 | 1.334 |
| **6DDF** | 0.930 | 0.916 | 0.869 | 1.019 | 0.859 | 1.111 | 1.027 | 1.193 | 1.056 | 1.152 |
| **6OS9** | 2.334 | 2.183 | 2.178 | 0.791 | 1.049 | 1.265 | 1.076 | 0.855 | 1.073 | 1.056 |

| [a] Corresponding LRMS [Å] | | | | | | | | | | |
|---|---|---|---|---|---|---|---|---|---|---|
| PDB ID | P1 | P2 | P3 | P4 | P5 | P6 | P7 | P8 | P9 | P10 |
| **3OE0** | 10.361 | 6.899 | 9.967 | 10.906 | 8.507 | 9.327 | 9.901 | 7.812 | 9.382 | 7.287 |
| **4GRV** | 3.275 | 2.783 | 4.109 | 1.987 | 2.997 | 2.577 | 3.485 | 2.827 | 4.767 | 2.447 |
| **5GLH** | 4.054 | 3.799 | 6.600 | 4.233 | 3.604 | 3.778 | 14.253 | 4.211 | 2.999 | 5.513 |
| **5XJM** | 3.760 | 4.624 | 3.541 | 5.212 | 4.643 | 3.894 | 4.967 | 4.188 | 4.629 | 4.128 |
| **6C1R** | 5.297 | 6.772 | 6.053 | 3.650 | 7.264 | 7.590 | 7.423 | 6.505 | 8.019 | 5.081 |
| **6DDF** | 3.003 | 2.891 | 2.902 | 3.446 | 2.530 | 4.035 | 3.467 | 4.297 | 3.575 | 4.295 |
| **6OS9** | 9.299 | 8.526 | 8.164 | 2.266 | 3.347 | 4.476 | 3.834 | 2.641 | 3.759 | 3.657 |

| [a] Corresponding Fnat values [Å] | | | | | | | | | | |
|---|---|---|---|---|---|---|---|---|---|---|
| PDB ID | P1 | P2 | P3 | P4 | P5 | P6 | P7 | P8 | P9 | P10 |
| **3OE0** | 0.207 | 0.310 | 0.241 | 0.133 | 0.180 | 0.103 | 0.172 | 0.310 | 0.213 | 0.155 |
| **4GRV** | 0.364 | 0.436 | 0.364 | 0.527 | 0.382 | 0.455 | 0.418 | 0.432 | 0.455 | 0.455 |
| **5GLH** | 0.435 | 0.361 | 0.231 | 0.269 | 0.343 | 0.324 | 0.000 | 0.352 | 0.417 | 0.211 |
| **5XJM** | 0.235 | 0.529 | 0.353 | 0.216 | 0.275 | 0.275 | 0.314 | 0.314 | 0.314 | 0.387 |
| **6C1R** | 0.292 | 0.208 | 0.292 | 0.479 | 0.312 | 0.396 | 0.208 | 0.292 | 0.333 | 0.354 |
| **6DDF** | 0.520 | 0.640 | 0.760 | 0.640 | 0.600 | 0.560 | 0.520 | 0.520 | 0.520 | 0.600 |
| **6OS9** | 0.180 | 0.220 | 0.320 | 0.440 | 0.360 | 0.280 | 0.420 | 0.440 | 0.440 | 0.460 |

[a]LRMS and Fnat values are calculated for the GPCR-peptide complex models presenting the lowest IRMS values.

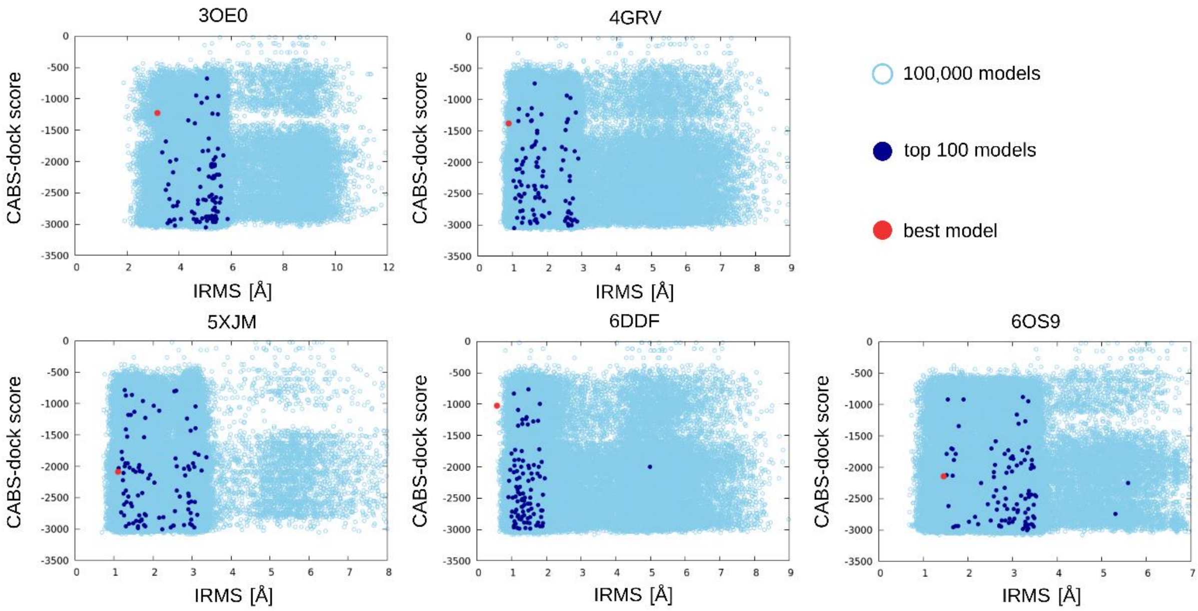


**Figure S1.** Plots presenting CABS-dock energy score versus IRMS [Å] values calculated for the models (PDB ID: 3OE0, 4GRV, 5XJM, 6DDF, 6OS9) generated during STAGE 1 of the modeling protocol (values calculated for models in CA representation). Plots correspond to data included in Table 1 of the manuscript.

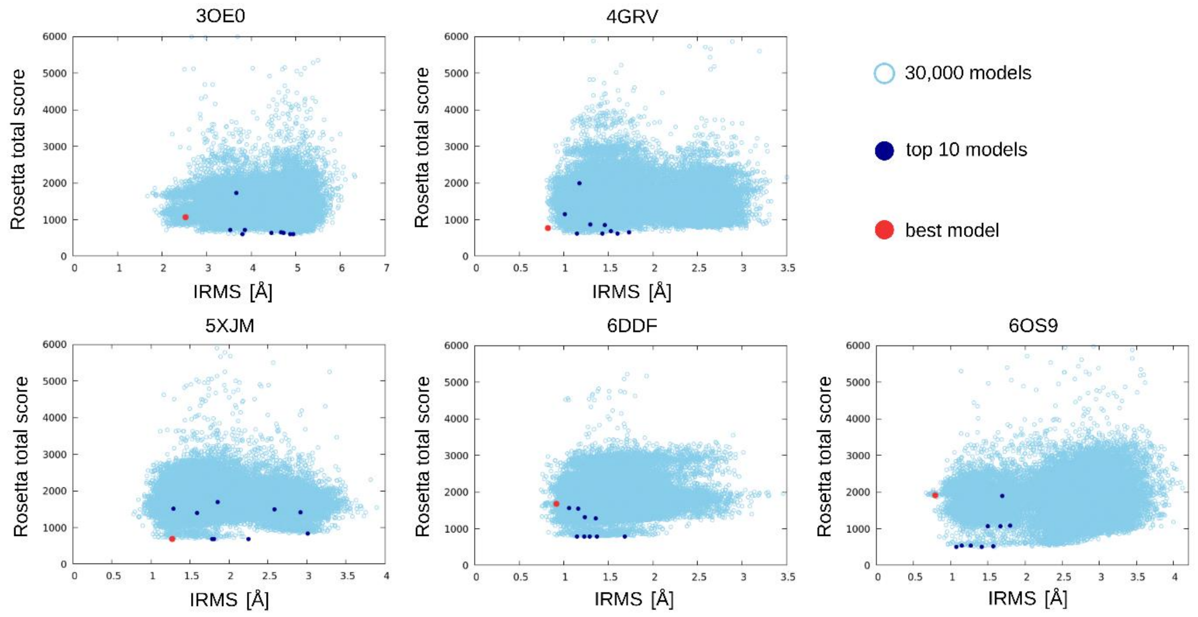


**Figure S2.** Plots presenting Rosetta FlexPepDock total score versus IRMS [Å] values calculated for the models (PDB ID: 3OE0, 4GRV, 5XJM, 6DDF, 6OS9) generated during STAGE 3 of the modeling protocol (values calculated for models in all-atom representation). Plots correspond to data included in Table 1 of the manuscript.

### SI References

1. Lensink MF, Velankar S, Wodak SJ, Modeling protein–protein and protein–peptide complexes: CAPRI 6th edition. Proteins: Structure, Function, and Bioinformatics, 2017. 85(3): 359-377.
